# Development of methods for purification of individual biological active substances obtained from extracts of *Hedysarum neglectum*

**DOI:** 10.1101/2021.06.06.447004

**Authors:** Anastasia Igorevna Dmitrieva, Margarita Yuryevna Drozdova, Vyacheslav Fedorovich Dolganyuk

**Affiliations:** Researcher at the Laboratory of Bioassay of Natural Nutraceuticals, Federal state budgetary educational institution of higher education Kemerovo State University; Laboratory assistant-researcher of the Laboratory of Bioassay of Natural Nutraceuticals, Federal state budgetary educational institution of higher education Kemerovo State University

## Abstract

*Hedysarum neglectum* is a forage plant. Xanthone glycoside - mangiferin is extracted from this plant and used for medicine “Alpizarin”. In addition to substances of xanthone nature (mangiferin and isomangiferin) Hedysarum neglectum contains sugars, vitamins and provitamins, tannins; in the underground part it contains oligomeric catechins, isoflavonoids, butylphenols, alkaloids, tannins, flavonoids, saponins, coumarins, carbohydrates, vitamin C. For selecting optimal schemes of fractionization of substances, it is necessary to resort to multi-stage schemes of group-wide (preliminary) isolation and preparative accumulation. In particular cases, it is necessary to take into account the presence of concomitant substances, as well as the effectiveness and selectivity of the sorption-chromatographic technologies used. According to the results of the studies, the use of silica gel and sefadex LH-20 for the isolation of a complex of flavonoids and gallic acid is the most effective method for the selection of the optimal variant of the preparative isolation of the total amount of BAS in mcg/ml. The results of the research allowed us to identify the target biologically active substances with a degree of extraction of at least 80%: - fractions of xanthones, flavonoids, gallic acid.

## Introduction

*Hedysarum neglectum* (Hedysarum gmelinii Ledeb.) is a forage plant [2]. From this plant. Xanthone glycoside - mangiferin is extracted from this plant, from which the medicine "Alpizarin", which has an antiviral effect, was produced [4].

In the work of S. A. Kubentaev, it is indicated that in addition to substances of xanthone nature (mangiferin and isomangiferin), *Hedysarum* in its ground part contains: sugars, vitamins and provitamins, tannins; in the underground part it contains oligomeric catechins, isoflavonoids, butylphenols, alkaloids, tannins, flavonoids, saponins, coumarins, carbohydrates, vitamin C [3].

Due to the content of such biological active substances (BAS), the plant has anti-inflammatory, antitumor, immunostimulating and tonic properties.

Plant cells and tissues, along with secondary metabolites of target substances (flavonoids, alkaloids, saponins, coumarins, etc.), contain products of general metabolism - primary metabolites: carbohydrates, organic acids, amino acids, proteins, etc., and also in significant quantities they may contain resinous substances, carotenoids, chlorophylls. Therefore, when choosing the optimal schemes of fractionation of substances, it is necessary to resort to multi-stage schemes of group-wide (preliminary) isolation and preparative accumulation. In particular cases, it is necessary to take into account the presence of concomitant substances, as well as the effectiveness and selectivity of the sorption-chromatographic technologies used.

The chromatographic mobility of compounds of plant origin is subject to strict laws and correlates with the chemical structure. Thus, the sorption-chromatographic parameters of one of the most common and practically significant group of biologically active compounds, flavonoids, are affected by such structural features as the variation in the number of hydroxyl groups in the molecule, the presence of methoxyl substituents, acithelination, glycosidation features (carbohydrate, the amount of sugar residues, their position), and, in fact, the ortho and vicial position of substituents.

## Objects and methods

– seeds of the *Hedysarum neglectum*, obtained and sprouted in the Botanical Garden of the Baltic Federal University named after I. Kant, Kaliningrad;
– alcoholic extracts of *Hedysarum neglectum*.

Ethyl alcohol was used as an extractant. The dried plant material was crushed in a mill of the brand LZM-1M (Russia, Olis) and sifted through a sieve with a hole size of 1 mm. The resulting raw materials were stored at room temperature in a dark room. Fine powder of the studied plant (3.0 g) was extracted in 260 ml of ethyl alcohol under static conditions to obtain BAS.

The extraction of plant material was carried out on a water bath of the PE-4310 brand (Russia, EKROSHIM) with a reverse refrigerator. The extraction frequency is 2, the concentration of the extractant is 70%, the extraction temperature is 30 °C, the extraction exposure is 6 h.

LC chromatography was used for the analysis of extracts of callus cultures of *Hedysarum neglectum*. The preparative isolation and accumulation of mangiferin was performed in the low-pressure chromatography mode using cross-linked agarose as a sorbent. Water-alcohol extracts were used in the examination. The extraction data after evaporation were chromatographed on CL-4 B sepharose gel (Pharmacia, Sweden) under working conditions. Gel volume was 10 ml, elution rate was 0.1-0.4 ml / min. The eluent was: deionized water and 0.01 M sodium hydroxide solution. Chromatography was performed on a BioLogic low-pressure chromatograph (BioRad).

A chromatographic column with a sorbent modified with octadodecyl and endcapped with aminophenol groups was used to identify the BAS of *Hedysarum neglectum*. As an eluent, a mixture of water: acetonitrile, 0.1% tributylamine 85:15 was used. Elution was performed in the isocratic mode, flow rate of 1 ml / min, elution time of 55 min. Detection was carried out using a fluorescent detector, the excitation wavelength was 350 nm, the detection wavelength was 415 nm, the collection of individual substances was carried out automatically using a fraction collector.

The selection of rational parameters for the extraction of BAS from plant extracts was primarily guided by the chemical composition, structure and properties of the target BAS and associated ballast compounds, which had to be separated from the target BAS [5, 6]. Also, when selecting rational parameters, a significant factor was the quantitative content of the target BAS in the analyzed sample, which had to be taken into account at the stages of concentration [1, 7].

Based on the above, the schemes of analysis, isolation, purification and preparative accumulation are unique in relation to a specific plant and the biologically active compound isolated from it, as well as individual in relation to the features of the technological process and the resulting extraction.

When developing methods for the purification of individual biologically active substances obtained from medicinal plant extracts, they were primarily guided by ensuring adequate purification from accompanying (ballast) components [8, 9].

## Results and discussion

The extraction of biologically active substances from the root crop extract of the *Hedysarum neglectum* includes the following steps:

1. The ethanol extract is dried in a vacuum on a rotary evaporator IKA RV 8 at a temperature of 450C.
2. The resulting residue is fractionated on a BioLogic low-pressure chromatograph (BioRad) using column chromatography on silica gel (qualifications for column chromatography, Sigma) using a mixture of petroleum ether - ethyl acetate as an eluent. Changing gradient of ethyl acetate 100: 0; 50: 1; 20: 1; 10: 1; 5: 1; 2: 1; 1: 1; and 0: 1.
3. Methanol is fed to the column for desorption of gallic acid.
4. The fractions are selected using a 300 ml fraction collector each. Thus, nine fractions are collected (fractions 1-9). From fraction 3, a crude mangiferin crystal is obtained, which is then recrystallized from the mixture: petroleum ether-acetone, in the ratio of 20:1.

**Figure 1.**
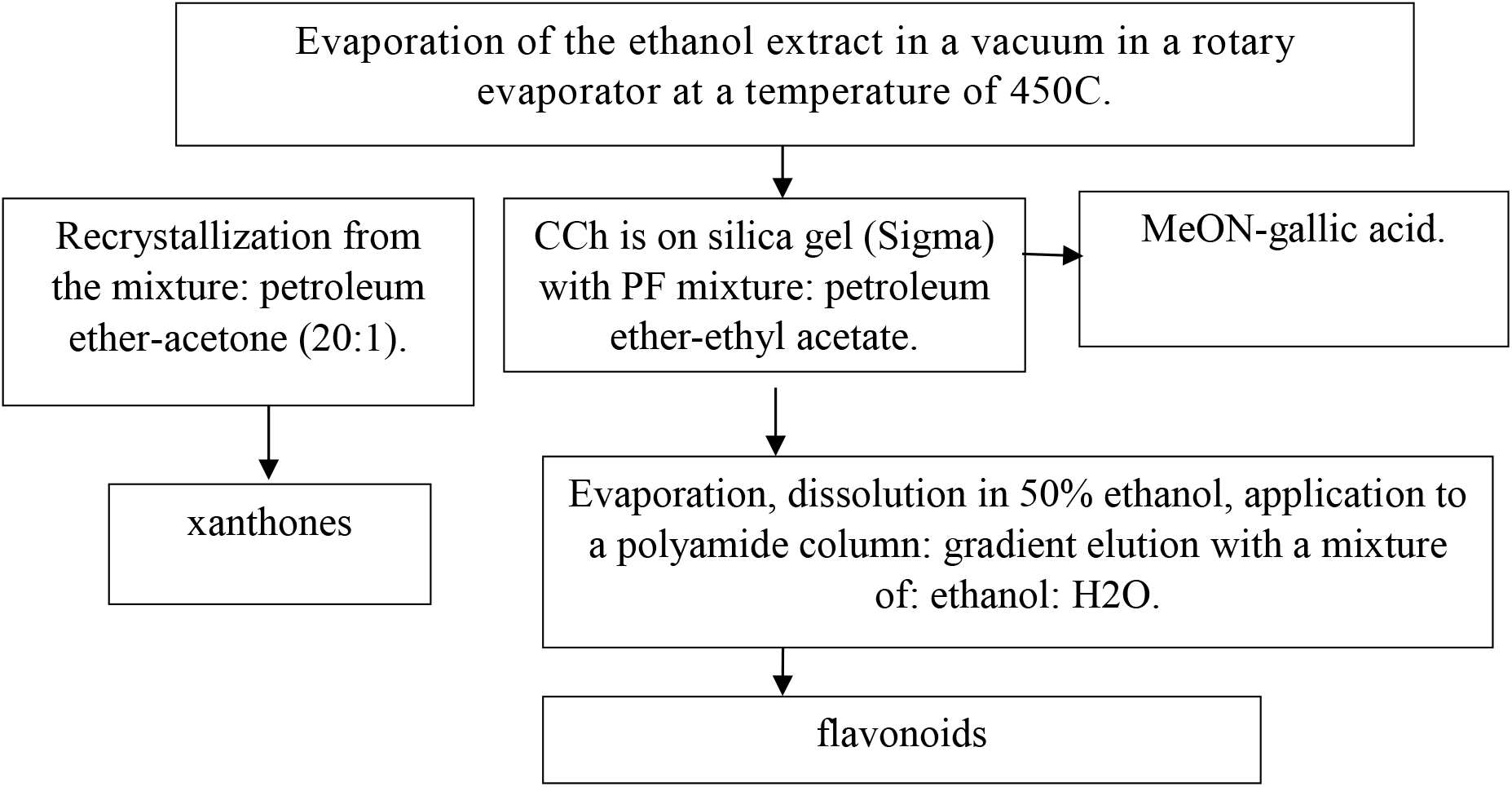
Scheme of isolation of biologically active substances from the extract of root cultures of *Hedysarum neglectum*.

When selecting rational parameters for the isolation of individual biologically active substances from extracts obtained from the biomass of callus, suspension cell cultures and root cultures of medicinal plants, various options for the preparative isolation of the total amount of biologically active substances were tested. The results of the studies are presented in table 1.

**Table 1.** Effectiveness of various variants of preparative isolation of the total amount of BAS

| Aluminum oxide | Silica Gel | Octodecylsilicagel<br>C18 | Cellulose | Sefadex<br>LH-20 | Activated carbon | Ion exchange resin | Organic solvents |
| --- | --- | --- | --- | --- | --- | --- | --- |
| 35,1<sup>FP</sup> | 24,1<sup>FP</sup> | 32,5<sup>FP</sup> | 14,31<sup>FP</sup> | 30,3<sup>FP</sup> | 35,3<sup>FP</sup> | 27,9<sup>FP</sup> | 35,26<sup>FP</sup> |
| ND | 80,8<sup>FL</sup> | 77,5<sup>FL</sup> | ND | 69,8<sup>FL</sup> | ND | ND | ND |
| ND | 57,02<sup>GA</sup> | 53,2<sup>GA</sup> | ND | 53,5<sup>GA</sup> | ND | 24,1<sup>GA</sup> | ND |

AR-Aromatic compounds (thymol, carvacrol)
GA-Gallic Acid
FL-Flavonoids
FP-Phenylpropanoids (oxycoric, coumaric acids)
\*n/a – not defined

When selecting rational parameters for the isolation of individual BAS from extracts of root cultures of *Hedysarum neglectum*, it was found that the use of this scheme for the isolation of biologically active substances allowed us to obtain fractions of xanthones, flavonoids, gallic acid, followed by the isolation of individual substances from the purified extract. According to the results of the studies presented in Table 1, the use of silica gel and sefadex LH-20 for the isolation of a complex of flavonoids and gallic acid is the most effective method for the selection of the optimal variant of the preparative isolation of the total amount of BAS in mcg/ml.

The results of the research allowed us to identify the target biologically active substances with a degree of extraction of at least 80%: - fractions of xanthones, flavonoids, gallic acid.

The method of purification of individual biologically active substances isolated from extracts of root cultures of *Hedysarum neglectum* includes the following steps:

1. The ethanol extract is dried in a vacuum on a rotary evaporator IKA RV 8 at a temperature of 450C. The resulting residue is fractionated on a BioLogic low-pressure chromatograph (BioRad) using column chromatography on silica gel and for column chromatography (Sigma) using a mixture of petroleum ether - ethyl acetate as an eluent. Changing gradient of ethyl acetate is in the following way: 100: 0; 50: 1; 20: 1; 10: 1; 5: 1; 2: 1; 1: 1; and 0: 1.
2. After that, methanol is fed to the column for desorption of gallic acid.
3. The fractions are selected using a collector of fractions of 300 ml each. Thus, nine fractions are collected (fractions 1-9). From fraction 3, a crude mangiferin crystal is obtained, which is then recrystallized from the mixture: petroleum ether-acetone, in a ratio of 20: 1, and purified by rechromatography on sepharose CL6B (Sigma - Aldrich) on a BioLogic low-pressure chromatograph (BioRad), which allows the pure substance of mangiferin to be isolated.
4. The total residue of fractions 1-9, except for fraction 3, is placed on a column with polyamide (Sigma) (BioLogic low-pressure chromatograph (BioRad)), where two fractions are isolated: 5-hydroxy-4-methoxy-8-prenyl-2-hydroxyisopropyl dihydrofurano-[4,5:6,7]-isoflavone and a fraction consisting of quercetin and quercetin-rhamnopyranoside.
5. Fractions containing 5-hydroxy-4-methoxy-8-prenyl-2-hydroxyisopropyl dihydrofurano-[4,5:6,7] - isoflavone and quercetin-rhamnopyranoside are purified by preparative TLC with sample application using an APA-2 automatic applicator. Conditions for the purification of substances:

a. the use of Sorbfil AF plates, a mobile phase consisting of a mixture of: petroleum ether-acetone, in a ratio of 8: 1, makes it possible to isolate 5-hydroxy-4-methoxy-8-prenyl-2-hydroxyisopropyl dihydrofurano - [4,5:6,7] - isoflavone;
b. the use of Sorbfil AF plates, a mobile phase consisting of a mixture of: petroleum ether-AcOEt, in a ratio of 5: 1, makes it possible to isolate quercetin-rhamnopyranoside.

Purification scheme of BAS is shown in figure 2.

The results of studies on the development of methods for individual purification of biologically active substances, allowed to identify the target biologically active substances with a purity of at least 95%: Gallic acid, pure substance mangiferin, 5-hydroxy-4-methoxy-8-prenyl-2-hydroxysteroiddehydrogenase-[4,5:6,7]-the isoflavone, quercetin-rhamnopyranoside.

**Figure 2.**
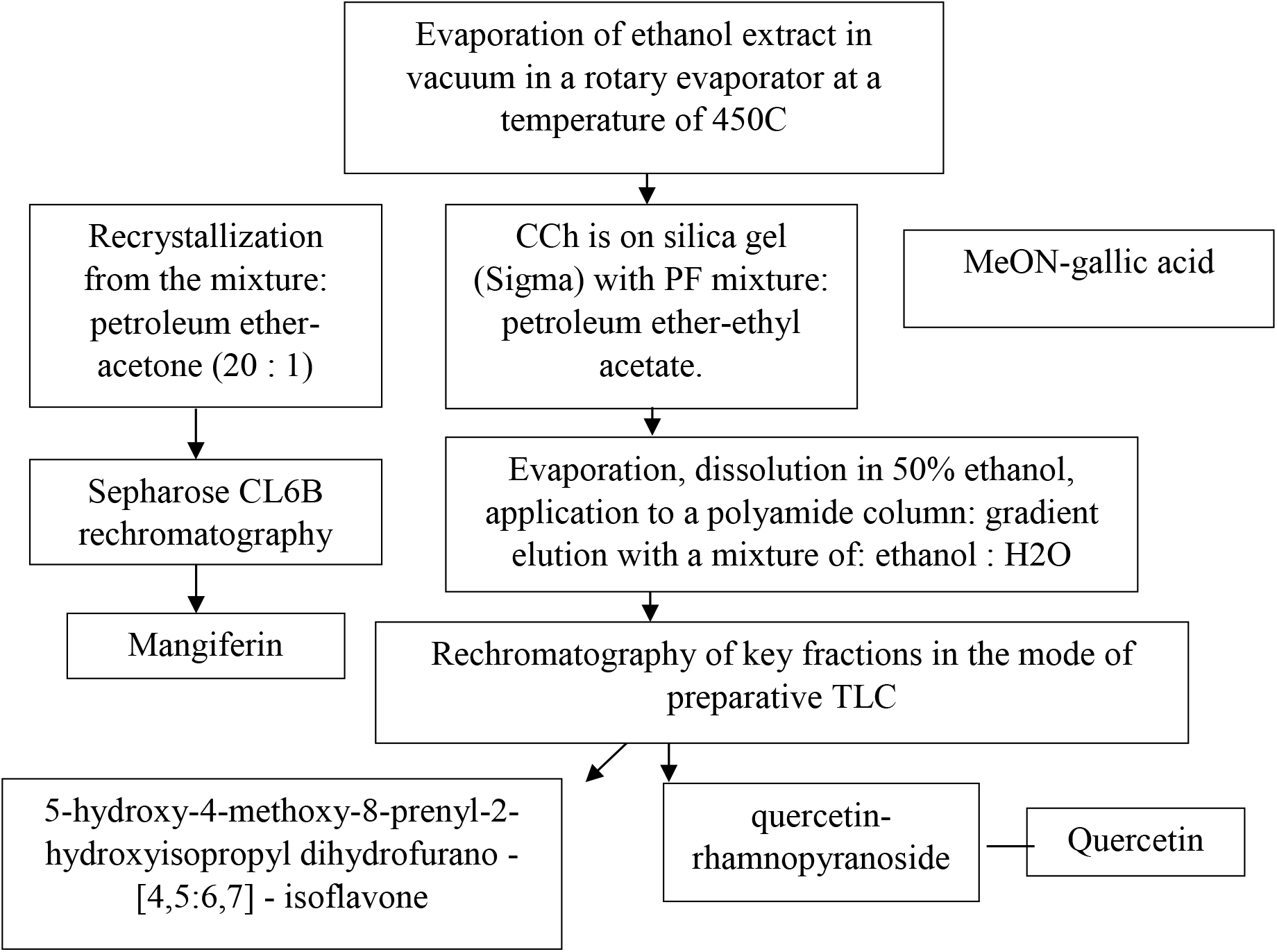
Scheme of purification of BAS obtained from the extract of root cultures of *Hedysarum neglectum*

**Figure 3.**
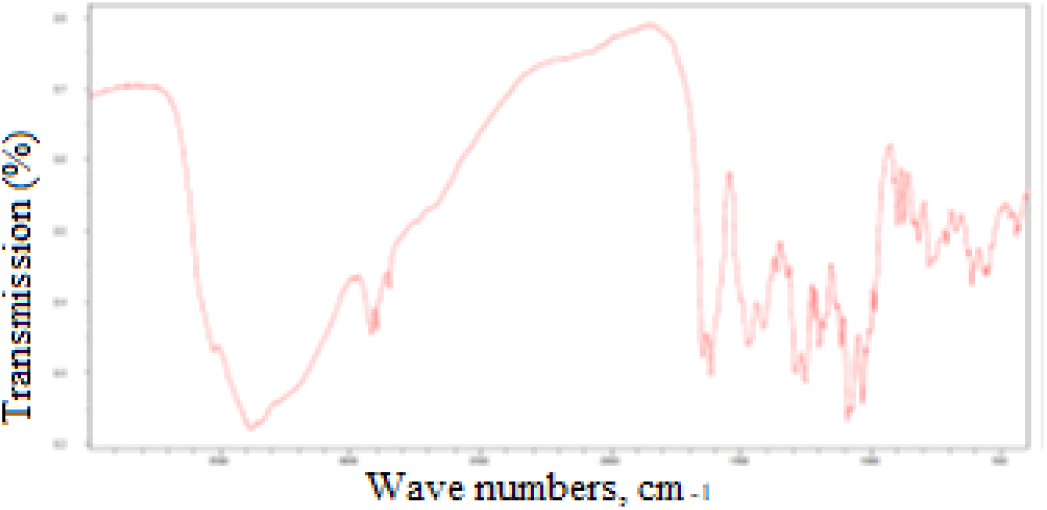
The IR spectrum of mangiferin

**Figure 4.**
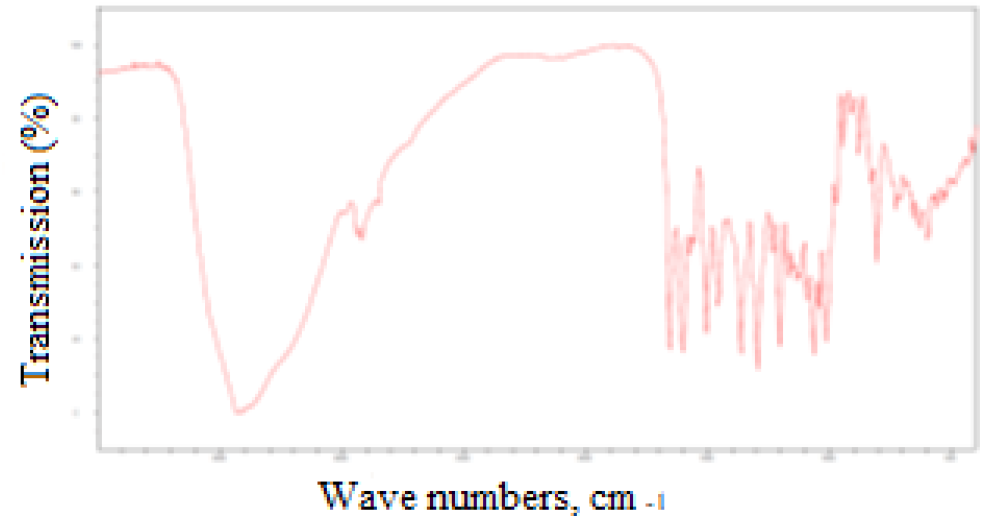
IR spectrum of quercetin

**Figure 5.**
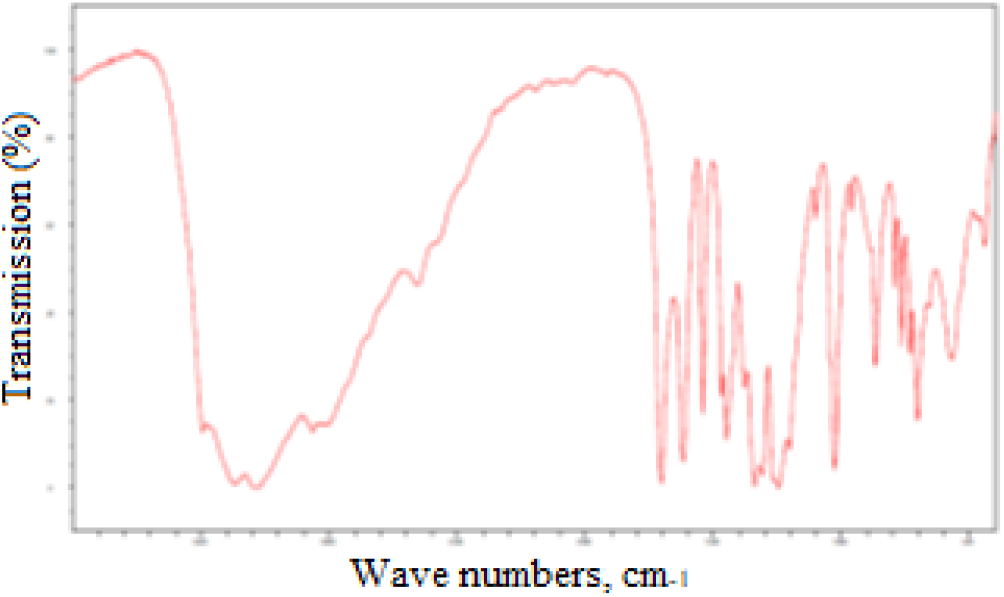
IR spectrum of gallic acid

**Figure 6.**
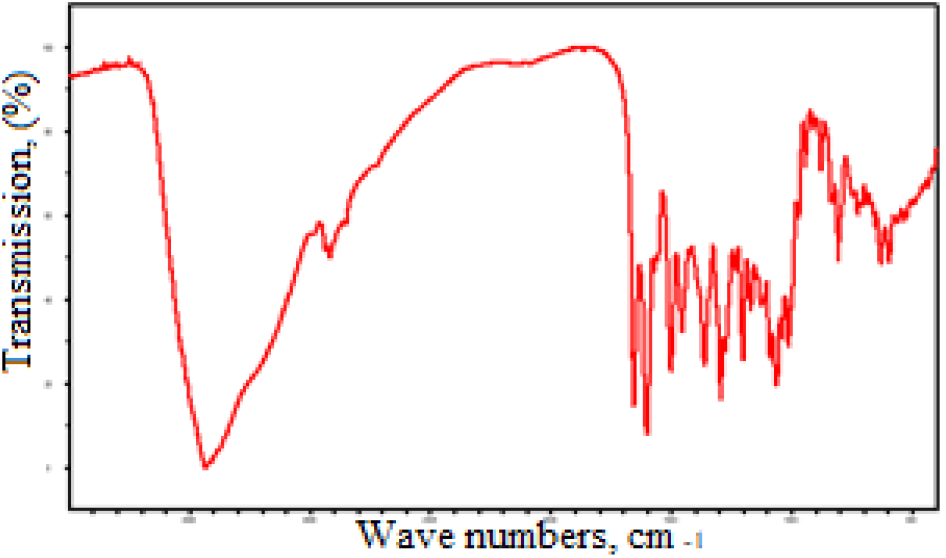
IR spectrum of quercetin-rhamnopyranoside

